# Variable latency between the founder genetic event and rhabdoid tumor expansion

**DOI:** 10.64898/2026.06.30.735306

**Authors:** Mònica Sánchez-Guixé, Alex Cebria-Xart, Naomi Fabre, Carlos J Rodriguez-Hernandez, Morena Pinheiro-Santin, Cinzia Lavarino, Jarno Drost, Ruben Van Boxtel, Núria López-Bigas, Alexandra Avgustinova, Abel González-Pérez

**Affiliations:** Institut de Recerca Sant Joan de Déu, Barcelona 08950, Spain; Institute for Research in Biomedicine (IRB Barcelona), The Barcelona Institute of Science and Technology, Baldiri Reixac, 10, 08028 Barcelona, Spain; Centro de Investigación Biomédica en Red en Cáncer (CIBERONC), Instituto de Salud Carlos III, Madrid, Spain; Princess Máxima Center for Pediatric Oncology, Utrecht, the Netherlands; Oncode Institute, Utrecht, the Netherlands; Division Cell Biology, Metabolism & Cancer, Department Biomolecular Health Sciences, Faculty of Veterinary Medicine, Utrecht University, Utrecht, The Netherlands; Universitat Pompeu Fabra, Barcelona, Spain; Institució Catalana de Recerca i Estudis Avançats (ICREA), Barcelona, Spain

## Abstract

Rhabdoid tumors are very aggressive rare pediatric cancers with poor survival affecting very young children. They are characterized by the bi-allelic loss of *SMARCB1* or *SMARCA4*, which is suspected to occur prenatally. However, their genomic evolution is not well understood. Here we assembled the largest cohort of whole-genome sequenced rhabdoid tumors to date, comprising 97 tumors from 88 children. We discovered that, in 42% of cases, the bi-allelic inactivation of the driver gene occurred via a Copy Number Neutral-Loss of Heterozygosity (CN-LOH). We exploited these CN-LOH events and the steady accumulation of age-related mutations in the tumor genomes to estimate the age of donors at the time of occurrence of the driver event and at the time of emergence of the clonal expansion. Across all cases with CN-LOH, the loss of the driver gene occurred very early during prenatal development. However, the clonal expansion that ultimately gave rise to the tumor occurred at different times during infancy, even several years after the acquisition of the founder event. These results indicate that probably other factors, besides the genetic driver event, are required to promote rhabdoid tumorigenesis.

## Introduction

Rhabdoid tumors are very aggressive childhood malignancies with devastatingly high mortality rates (<20% of 4-year survival)^1,2^. They affect mostly very young children (0-5 years old), although in some rare cases they affect children at puberty^1^. Rhabdoid tumors can appear throughout the body, and are mainly classified as extra-cranial –also called Malignant Rhabdoid Tumors, MRTs– or intra-cranial –also known as Atypical Teratoid/Rhabdoid Tumors, ATRTs. MRTs are detected frequently in the kidneys, liver, and soft tissues. Although considered rare, MRTs account for 9-18% of all extra-cranial solid malignancies in children younger than one year old^1^, and ATRTs account for 10-15% of all malignant brain tumors in children under the age of three^3^.

Rhabdoid tumors are genomically very simple, usually driven by the bi-allelic loss of *SMARCB1*, or, in rare cases, *SMARCA4*^1^. *SMARCB1* and *SMARCA4* encode, respectively, a core subunit and an ATPase of the SWI/SNF chromatin-remodeling complex that plays critical roles in epigenetic and transcriptional regulation^4^. Furthermore, *SMARCA4* mutated tumors represent a separate molecular subtype entity different from the subtypes identified in *SMARCB1* mutated tumors^5^. Apart from *SMARCB1* and *SMARCA4* mutations, rhabdoid tumors exhibit low mutational burden, and in general have diploid genomes^6,7^.

Despite their simple genomic make-up, the precise origin of rhabdoid tumors during prenatal development and their evolution is not clear. Previous studies in mouse models^8,9^ and the phylogenetic analysis of two patient cases^10^ point to a prenatal origin of the driver event. However, the complete genomic history of these tumors is poorly understood. Specifically, whether the loss of *SMARCB1* or *SMARCA4* alone is sufficient for their oncogenic transformation and tumor outgrowth, or whether other potentially non-genetic events are required to promote these processes (as suggested in a previous study that found normal cells with the *SMARCB1* null genotype in a patient suffering from a rhabdoid tumor^10^), remains unresolved.

In this study, we collected the largest cohort of whole-genome sequenced rhabdoid tumors to date (88 cases), and carried out the first systematic analysis of the genomic evolution of these tumors. We discovered that in almost half of the cases (42%), a Copy Number Neutral-Loss of Heterozygosity (CN-LOH) was responsible for the loss of the second allele of the driver tumor suppressor gene. Exploiting age-related somatic mutations overlapping the duplicated genomic region, we confirmed that, across all rhabdoid cases, this CN-LOH event occurred very early in life, very likely during prenatal development, irrespective of the age of the patients at diagnosis. We also found that the clonal expansion from the most recent common ancestor of the rhabdoid tumors occurred at different ages across cases, in general relatively shortly before the diagnosis of the malignancy. This means that the time between the occurrence of the genetic driver event and the beginning of the clonal expansion of the rhabdoid tumors varied between a few months and several years. This finding suggests that an event or a likely non-genetic event or process is necessary for the development of the disease after the acquisition of the driver event.

## Results

### Genomic and clinical overview of 88 cases diagnosed with rhabdoid tumor

We assembled a large cohort of whole-genome sequenced rhabdoid tumors. This comprised 98 samples obtained from 89 children. Sixteen of these cases (22 tumors; 18%) were recruited at *Sant Joan de Déu Barcelona Children’s Hospital* (SJD) and sequenced (together with matched normal blood samples) in-house. The genomic sequence of 4 tumors from a case (1,1%) and its matched normal blood sample was obtained from *Princess Maxima Center*, the Netherlands (PMC). The tumor and normal genomic sequence of 56 cases (62,9%) from *Therapeutically Applicable Research to Generate Effective Treatments* (TARGET) program^7,11^, and 16 cases (18%) from *St Jude Cloud* (StJude)^12^ were also obtained. We then identified all somatic genomic variants (single nucleotide variants, or SNVs, small insertions and deletions, or indels, structural variants, or SVs and Copy Number Alterations, or CNAs) (see **Methods**–*Analysis of driver mutations)*. Of these 89 cases, 20 (22.5%) were found in the central nervous system, and therefore classified as atypical teratoid rhabdoid tumors (ATRTs); the remaining 69 (77.5%) were extra-cranial malignant rhabdoid tumors (MRTs), of which 53 (59.6%) were located in the kidney. Most kidney-related MRTs (56) were in the TARGET cohort, while most (15) ATRTs were included in the StJude cohort (**Extended Data Fig. 1a**, **b** and **c**). Forty-three patients (48%) were female and forty-six (52%) were male (**Fig. 1a**). The age at diagnosis in most cases is between 0 and 2 years old, although some cases were diagnosed at older ages, up to 15 years (**Extended Data Fig. 1d**).

**Fig. 1:**
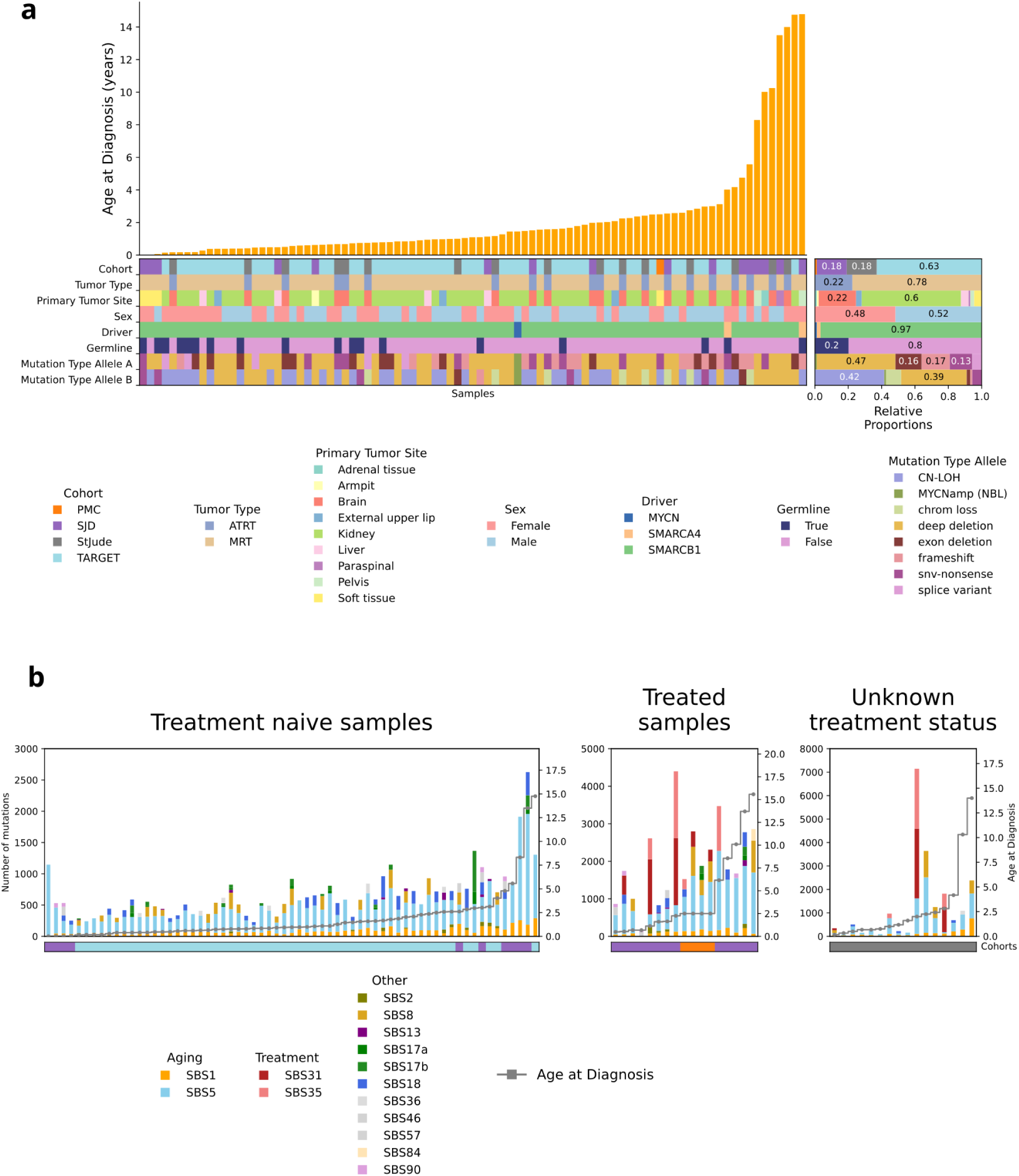
Genomic and clinical overview of 88 cases with rhabdoid tumors. **a**, Clinical information overview of the collected cohort. The box on the right accounts for the relative proportions of each category. Take note that *MYCN*amp sample from TARGET project is included, which is later identified as a neuroblastoma instead of a rhabdoid tumor (see **Supplementary Methods**–*Mislabeled neuroblastoma sample*); **b**, The landscape of mutational signatures in rhabdoid tumors. Tumor mutational burden per sample in increasing age at diagnosis from left to right (age is indicated in grey), each color represents the fraction of mutations corresponding to the given mutational signature.

We identified a loss-of-function alteration affecting both alleles of the *SMARCB1* gene in 86 (96.6%) of the cases. In 2 cases, a loss-of-function alteration affecting the two alleles of *SMARCA4* was found (**Supplementary Table 1**, **Fig. 1a, Extended Data Fig. 2**). The remaining case was found to harbor a *MYCN* amplification and was therefore considered an incorrectly classified neuroblastoma, and excluded from the study (**Extended Data Fig. 3a** and **b**, see **Supplementary Methods**–*Mislabeled neuroblastoma sample*). In 18 cases (20.2%), one allele of the driver gene was affected by a germline variant. Intriguingly, we found a case with two germline alterations affecting *SMARCB1* (PT76 of the TARGET cohort^7^) (**Extended Data Fig. 3 c**, **d** and **e**, see **Supplementary Note**–*A case with two germline SMARCB1 variants*).

In forty-two cases (47.2%), the loss of one of the alleles of the driver gene was caused by a large deletion (i.e., involving the whole gene, and in some instances, a big part of the chromosome). In 30 (33.7%) of these cases, the second allele was also inactivated by a large deletion, while in eight (9%) it was affected by a loss of one copy of chromosome 22. In most cases affected by chromosome 22 large deletions, these involve a large part of the chromosome, commonly affecting from *SMARCB1* position to the end of the q arm (**Extended Data Fig. 2c**). Smaller variants caused the inactivation of one allele of the driver gene in 49 cases, including exon deletions (affecting mostly exons 1 and 6) in 16 cases (18%), frameshift indels in 16 cases (18%), nonsense small nucleotide variants (SNVs) in 17 cases (19%), and splice-affecting variants in 5 cases (6%) (**Fig. 1a, Extended Data Fig. 2b**, **c**, and **d**). Interestingly, in 37 cases (42%), a Copy Number Neutral-Loss of Heterozygosity (CN-LOH) caused the loss of the second allele of the driver gene.

We then analysed the mutational processes active in the samples. We observed that most mutations in treatment-naive samples could be attributed to clock-like mutational processes (represented by SBS1 and SBS5), while several cases also showed activity of SBS18 (potentially linked to oxidative damage) and SBS8, a mutational signature of unknown etiology (**Fig. 1b**). We also observed one case of the TARGET cohort (PT76, carrying two germline *SMARCB1* mutations; see above^7^), and another of the SJD cohort (represented by samples PT86-T1 and PT86-T2) with activity of SBS17a and SBS17b, previously reported in tumors arising from organs of the digestive tract^13,14^ (**Extended Data Fig. 4a**, see **Supplementary Note**–*A case with two germline SMARCB1 variants*). Across tumors obtained from treatment-exposed patients, we identified platinum-related mutational signatures (SBS31 and SBS35). These signatures also appeared across 4 tumors obtained from patients in the StJude cohort, with unknown treatment data (**Fig. 1b**).

### The most recent clonal expansion of rhabdoid tumors occurs at different ages

Somatic SNVs (mutations, for simplicity) that accumulate steadily throughout life (e.g., those contributed by SBS1 and SBS5) can be used as a clock to estimate the time elapsed between fertilization and the moment at which the most recent common ancestor (MRCA) cell of the rhabdoid tumor began its clonal expansion. All age-related mutations of this rhabdoid MRCA will be present in all the cells of the tumor (clonal mutations) (**Fig. 2a**).

**Fig. 2:**
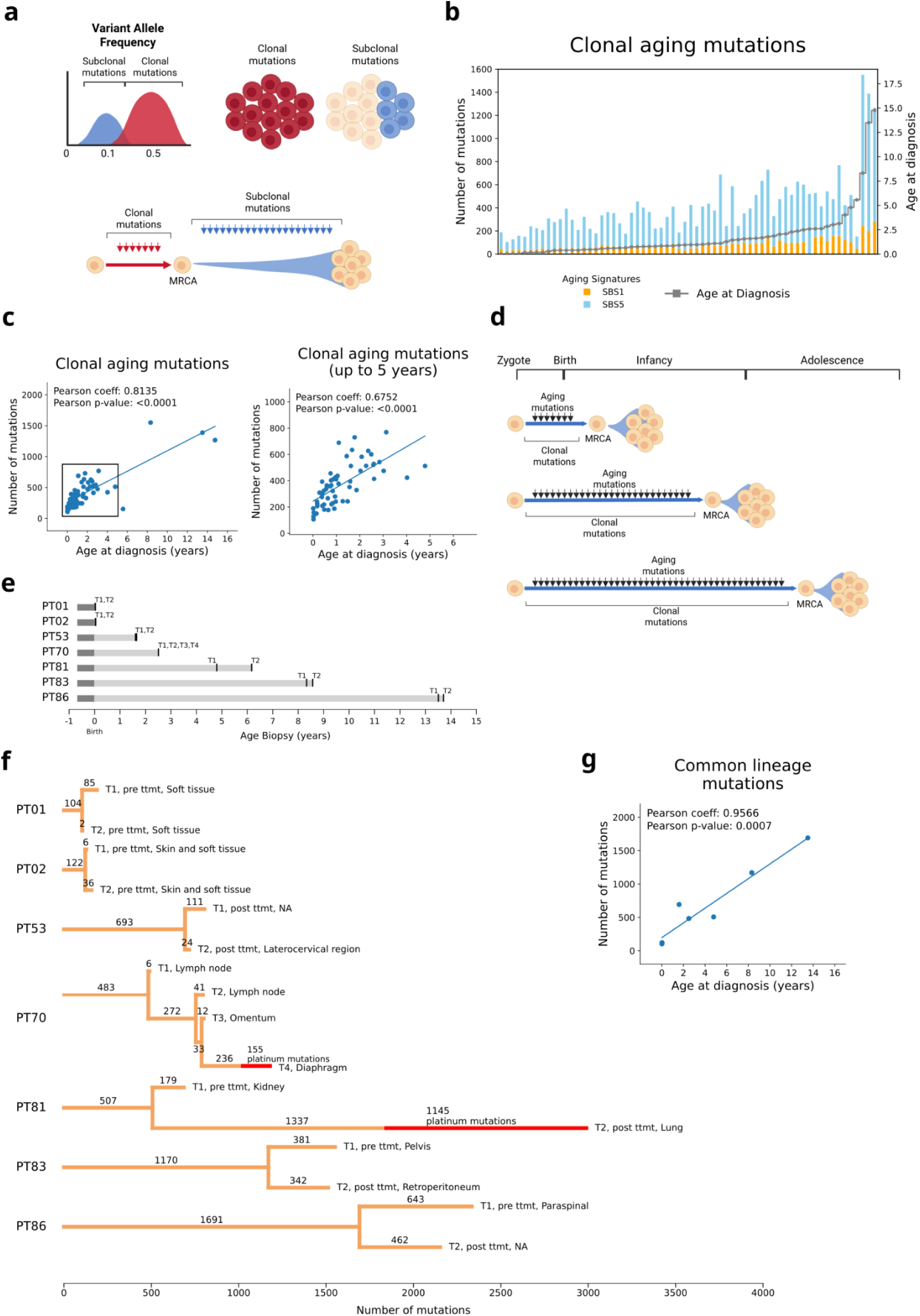
Relative timing of the clonal expansion of the Most Recent Common Ancestor (MRCA) **a**, The variant allele frequency distribution allows for the separation of clonal and subclonal mutations, where clonal mutations are those belonging to all the sequenced cells and subclonal mutations those belonging to a subset of cells. Clonal mutations are those that harbored the MRCA; **b**, Clonal aging mutations (signatures SBS1 and SBS5) of treatment naive samples, ordered by age at diagnosis (from younger–left– to older –right–, indicated in grey); **c**, Left panel: correlation of the clonal aging mutations with the age at diagnosis per treatment naive sample (Pearson’s coefficient: 0.8135, p-value < 0.0001); right panel: Correlation of the clonal aging mutations with the age at diagnosis per treatment naive sample in a selection of cases with up to 5 years old (Pearson’s coefficient: 0.6752, p-value < 0.0001); **d**, Representation of the clonal expansions at different times across the different cases: low number of clonal aging mutations indicates that the tumor expanded earlier and higher number of clonal aging mutations indicates that the tumor expanded later; **e**, Age at biopsy of each sample for those cases with two or more tumor samples available; **f**, Tumor evolutionary trajectories of cases with two or more samples. The length of the bars are proportional to the number of mutations (indicated); corresponding platinum mutations are shown in red; **g**, Correlation of the number of mutations in the common lineages with the age at diagnosis (Pearson’s coefficient: 0.9566, p-value = 0.0007).

Counting these age-related clonal mutations in the tumors in this cohort, we next aimed to discern whether the clonal expansion of the rhabdoid MRCA starts at approximately the same age across patients, independently of their age at diagnosis, or whether it is initiated at different ages. If the first hypothesis were true, we expected to observe approximately the same number of age-related clonal mutations across treatment-naive tumors (N=64; excluding the tumor with two *SMARCB1* germline variants; **Extended Data Fig. 3c**, **d** and **e**), irrespective of the age at diagnosis.

However, contrary to this postulate, we observed very different numbers of age-related clonal mutations across treatment-naive rhabdoid tumors (**Fig. 2b**, **Extended Data Fig. 4b**). Indeed, the number of these mutations showed significant correlation with the age of the patients at diagnosis (Pearson’s coefficient = 0.8135 and p-value < 0.0001, **Fig. 2c**, left panel). We obtained the same result when the analysis was restricted to patients diagnosed at 5 years old or younger (Pearson’s coefficient = 0.6752 and p-value < 0.0001, **Fig. 2c**, right panel). This result indicates that the most recent clonal expansion that gave rise to the rhabdoid tumor occurred at very different ages across patients; the later the start of these rhabdoid clonal expansion, the later the age at diagnosis (**Fig. 2d**).

### Recent divergence of multiple rhabdoid lineages

Studying the cases with two or more tumors, we next sought to distinguish whether the divergence between different sequenced tumors was relatively older or more recent (with respect to the moment of diagnosis) (**Fig. 2e**). To this end, we split the clonal mutations identified in each tumor into those shared with other tumor(s) (i.e., occurred before lineage divergence), and those unique (**Extended Data Fig. 5a**). As expected, in all cases the driver mutation is one of the shared events. In cases PT01, PT02, PT53, PT70, PT83 and PT86, most clonal mutations in the tumors were shared across different tumors, indicating a relatively recent time of divergence between them. Only case PT81 displays signs of separate evolution of the two tumor samples, a primary lesion in the kidney detected at 4 years and 9 months of age and a relapse after the treatment of the primary tumor in the lung at 7 years and 1 month of age. Since platinum-related mutations in the relapse tumor are clonal, we can deduce that this relapse emerged from a full clonal expansion that occurred after the treatment, starting from a single cell of the original rhabdoid tumor^15^. This probably explains the extended time of evolution of the relapse lesion, which expanded from a single cell of the primary tumor (**Extended Data Fig. 5b**, see **Supplementary Note**–*Evolution of multiple rhabdoid tumors in an individual*). In case PT70, with four separate rhabdoid tumor lesions (T1-4) collected at the autopsy, we observe three tumors with platinum mutations. In T2 and T3, all platinum-related mutations were subclonal, indicating that their clonal expansions started before the exposure to the treatment. However, in T4, a combination of clonal and subclonal platinum-related mutations indicated that the clonal expansion started during the exposure to treatment (**Extended Data Fig. 5c**, see **Supplementary Note**–*Evolution of multiple rhabdoid tumors in an individual*). In summary, a recent (with respect to diagnosis) and rapid process of divergent evolution without any obvious gain of new driver events occurs across rhabdoid cases. The bottleneck created by the treatment can perturb this process of divergence. Overall, the length of the common lineages also correlates with the age at diagnosis (Pearson’s coefficient=0.9566 and p-value=0.0007, **Fig. 2g**), indicating that all tumor samples analysed expanded relatively close to the age of diagnosis.

### Relative time of the driver gene bi-allelic loss across rhabdoid cases

We exploited the 37 cases (42%) of the cohort with a CN-LOH event implicating *SMARCB1* or *SMARCA4* to determine the relative time at which it occurs (from the zygote to the MRCA) across cases (**Fig. 3a, b**). In 35 cases, the CN-LOH event affected chromosome 22 (involving *SMARCB1* gene) and in the other 2, chromosome 19 (involving *SMARCA4* gene). The CN-LOH events affecting chromosome 22 usually cover the region from the genomic position of *SMARCB1* to the end of the q arm. In the case of chromosome 19, they cover the region between *SMARCA4* and the end of the p arm (**Extended Data Fig. 6**).

**Fig. 3:**
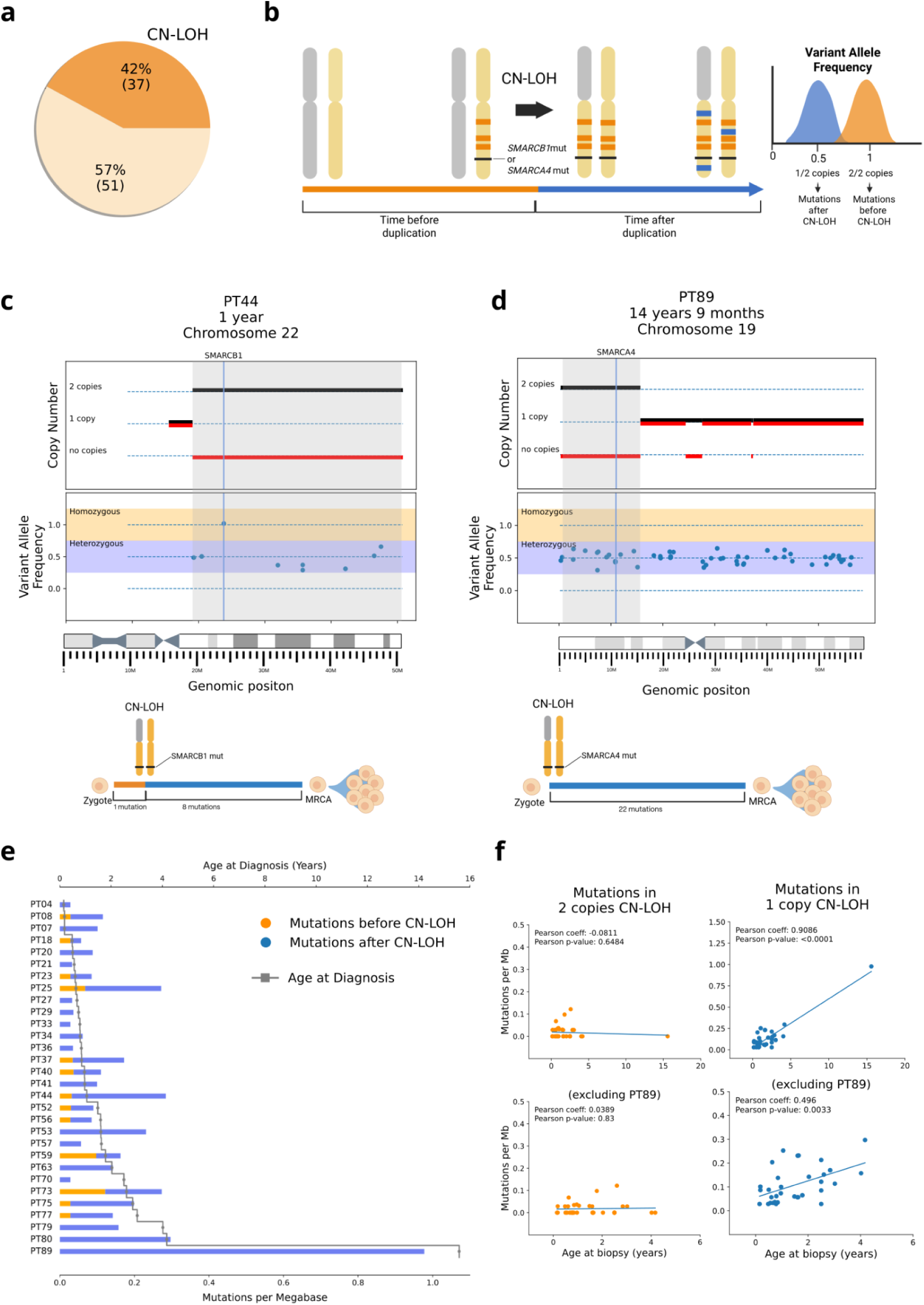
Relative timing of the founder event using the duplicated area of the Copy Number Neutral-Loss of Heterozygosity (CN-LOH). **a**, A fraction of 42% (37 cases) of the collected cohort harbors a CN-LOH; **b** Toy representation of a CN-LOH, showing the variant allele frequency distribution of the mutations in the duplicated area of the CN-LOH; orange mutations occur before the duplication and are therefore duplicated, and shown at a variant allele frequency of 1; blue mutations occur after the duplication and are unique to each chromosomic copy, and shown at a variant allele frequency of 0.5; **c** and **d**, CN-LOH in two representative cases; the grey area represents de CN-LOH range, the orange area represents the duplicated mutations (before the duplication, homozygous), and the blue area represents the unique mutations (after the duplication, heterozygous); all mutations shown are clonal; panels below are a toy representation of the relative timing of the *SMARCB1* and *SMARCA4* CN-LOH, where orange bars indicate the number of duplicated mutations and blue bars indicate the number of unique mutations; **c**, Case PT44, CN-LOH in chromosome 22; blue line indicates the position of *SMARCB1* gene; **d**, Case PT89 CN-LOH in chromosome 19; blue line indicates the position of *SMARCA4* gene; **e**, Relative timing of the CN-LOH across all cases with CN-LOH in the collected cohort; orange bars represent the number of mutations per megabase in homozygosity in the CN-LOH area (before duplication), and blue bars indicate the number of mutations per megabase in heterozygosity in the CN-LOH area (after duplication); samples are ordered from younger ages (top) to older ages (bottom); age at diagnostis is indicated with the grey line; **f**, Upper left panel: correlation between the number of duplicated mutations (before CN-LOH) per megabase in the CN-LOH with the age at diagnosis (Pearson’s correlation = −0.0811, p-value = 0.6484); Upper right panel: Correlation between the number of unique mutations (after CN-LOH) per megabase in the CN-LOH with the age at diagnosis (Pearson’s correlation = 0.9086, p-value < 0.0001); Lower panels: same correlations excluding PT89 sample with very high age at diagnosis (mutations in 2 copies CN-LOH, Pearson’s correlation = 0.0389, p-value = 0.83; mutations in 1 copy CN-LOH, Pearson’s correlation = 0.496, p-value = 0.0033).

We reasoned that age-related mutations accumulated in the genomic region affected by the CN-LOH before the duplication event will appear in both copies of the duplicated genomic region (homozygous; orange in **Fig. 3b**), while those accumulated after the event will appear in only one copy (heterozygous; blue in **Fig. 3b**). It is possible to distinguish these two sets of mutations by their variant allele frequency (VAF). Those acquired prior to the duplication (homozygous) will have double VAF (**Fig. 3b**). By counting the number of age-related mutations accumulated by the time of duplication, we can infer the relative time at which the duplication occurred. For instance, in PT44, we observe a CN-LOH in chromosome 22 (grey area, **Fig. 3c**). In this case, only 1 homozygous SNV is observed, which is in fact an intronic variant in *SMARCB1* (orange) –different from the driver mutation, which is a frameshift indel. All other SNVs overlapping the duplicated area are heterozygous (8 mutations, unique to each copy, in blue). In a second example, PT89, we observed a CN-LOH in chromosome 19, affecting *SMARCA4* (**Fig. 3d**). In this case, there is no somatic SNV in homozygosity (the first *SMARCA4* allele is affected by a germline variant), but there are 22 heterozygous mutations (unique to each copy, blue and gray area), indicating that a longer period of time has elapsed between the CN-LOH and the start of the clonal expansion of the rhabdoid MRCA than in the previous case.

Across 34 cases (after excluding 2 cases with tumor purity below 0.5 and a case with treatment-related clonal mutations) carrying a CN-LOH driver event we found fewer mutations occurring before the CN-LOH (orange) than after the CN-LOH (blue), indicating that this event took place closer in time to the first division of the zygote than to the clonal expansion of the rhabdoid MRCA (**Fig. 3e**, **Extended Data Fig. 7**). Moreover, the number of mutations accumulated before the CN-LOH (orange) did not show a significant correlation with the age of patients at diagnosis (Pearson’s coefficient = −0.0811, p-value = 0.6484) (**Fig. 3f**, upper left panel), not even after excluding a PT89 case which has high age at diagnosis (Pearson’s coefficient = 0.0389, p-value = 0.83) (**Fig. 3f**, lower left panel). Conversely, mutations accumulated after the CN-LOH (blue) showed a significant correlation with the age at diagnosis (Pearson’s coefficient = 0.9086, p-value < 0.0001) (**Fig. 3f**, upper right panel), even after excluding PT89 (Pearson’s coefficient = 0.496, p-value = 0.0033) (**Fig. 3f**, lower right panel). This indicates that the age at which the founder genomic event of the tumor –the bi-allelic loss of *SMARCB1* or *SMARCA4*– is acquired is not related to the age at diagnosis. In other words, the age at which the founder event is acquired does not appear to influence the speed at which the tumor evolves.

### The bi-allelic inactivation of the driver gene likely occurs during prenatal development

We next reasoned that the rate of accumulation of clonal age-related mutations could be used to convert the number of mutations accumulated before the CN-LOH to compute the chronological age at which this event occurred. To compute the yearly rate of accumulation of mutations, we compared the number of age-related clonal mutations identified across the whole genome of 64 treatment-naive tumors in the cohort with the embryonic age (that is, starting nine months prior to birth to accommodate mutations acquired during embryonic development) of the patients at diagnosis. We fitted the relationship between these two variables with two linear regressions, accounting for different prenatal and extra-uterine life mutation rates (indicated by significantly higher mutation rate in tumors of children diagnosed between birth and 2 years of age; **Fig. 4a** and **b; Extended Data Fig. 8a**^16–18^, see **Methods**–*Mutation rate calculation*). Importantly, the use of clonal mutations (that is, ignoring mutations that have accumulated in tumor cells after the last clonal sweep of the malignancy) in this calculation yields an underestimated mutation rate (**Fig. 4c, Extended Data Fig. 8b**).

**Fig. 4:**
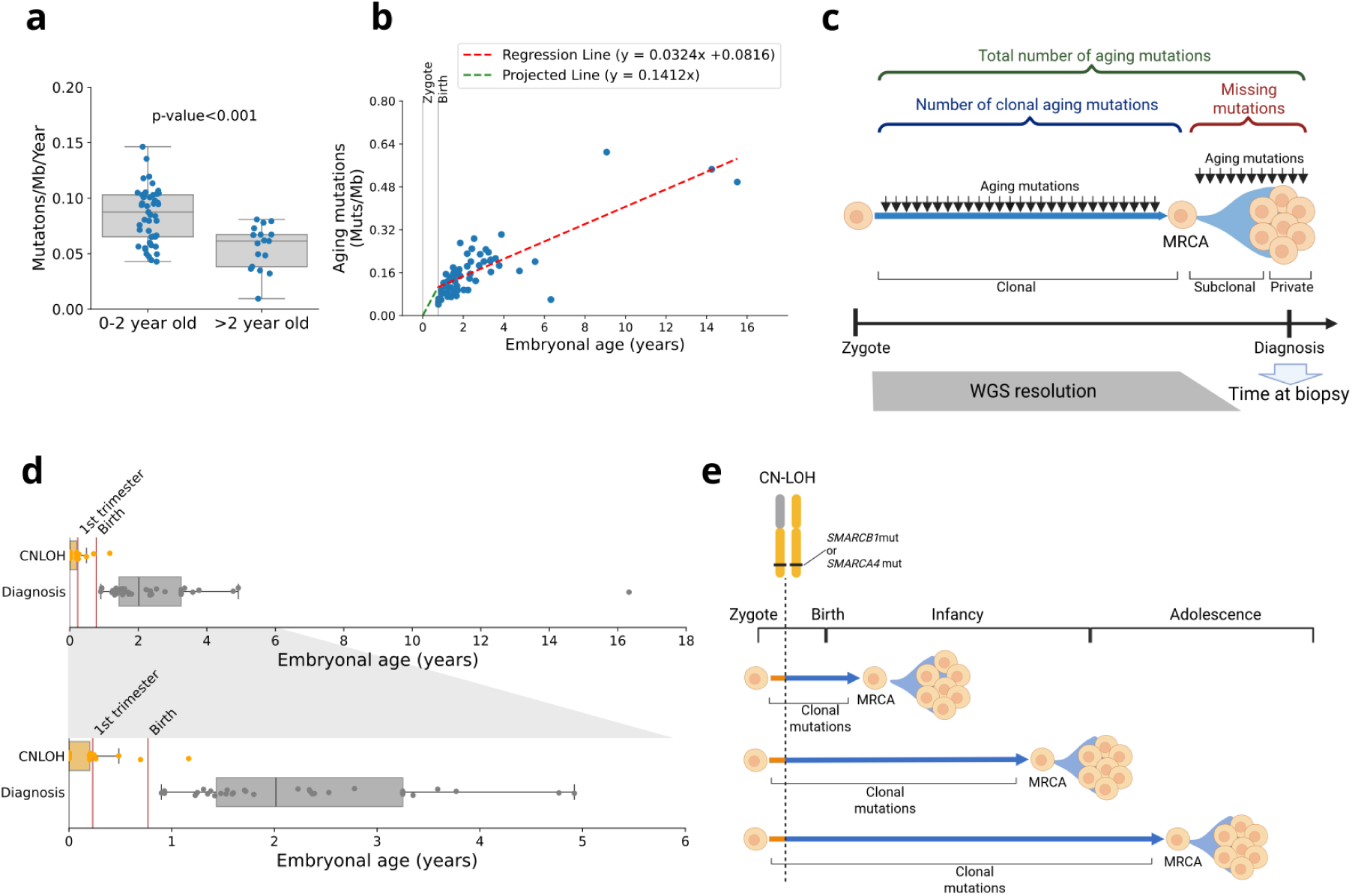
Very early occurrence of the driver founder mutation. **a**, Difference between mutation rate of cases with younger age at diagnosis (0-2 years old) and older age at diagnosis (>2 years old); Mann-Whitney U test, p-value < 0.0001; **b**, Mutation rate model with two slopes; clonal aging mutations of all treatment naive cases are used to plot a regression line from 9 months (birth) onwards (red slope), and another slope is projected from time 0 (zygote) to 9 months (green slope); **c**, Toy representation of the mutation rate approximation using the clonal mutations; regular bulk whole genome sequencing has a resolution limit, being able to identify clonal mutations and some subclonal mutations, but missing private mutations and many subclonal mutations; a mutation rate calculated with the clonal mutations (blue) and the time at biopsy will always have a gap of N mutations occurring from the clonal expansion to the time at biopsy (red); **d**, Aggregated estimated timings across all cases with CN-LOH of the CN-LOH timing (orange); the age at diagnosis is represented in grey; panel at the bottom represents a magnification of the data from 0 to 6 embryonic years (diagnosis age + 9 months of pregnancy); **e**, Toy representation of the very early timing occurrence of the CN-LOH (dashed vertical line) and the timing occurrence of the MRCA expansion.

We used this two-line model fitted on the mutation rate of tumors to calculate the age of occurrence of the CN-LOH in each case (**Extended Data Fig. 9**). The calculation consists of interpolating the density of SNVs (mutations per megabase) prior to the event into the regression model to obtain the time that explains the accumulation of this mutation density. The results show that in most of the 34 cases, the CN-LOH event was very likely acquired during prenatal development (**Fig. 4d**), in agreement with the previous hypotheses^9,10,19^. Only in one case could the calculation be consistent with a CN-LOH after birth. Nevertheless, since the mutation rate is underestimated in our calculation (that is, lower-than-real accumulated mutations per megabase per year), the real time of occurrence of the duplication event is overestimated, as probably less time than calculated can explain the observed number of homozygous SNVs. These results indicate that the founder event of rhabdoid tumors is very likely to occur during prenatal development. In any case, there does not appear to be any relationship between the age at which this event occurs and the age of diagnosis of the tumor. Conversely, the MRCA clonal expansion seems to occur relatively close to the age of diagnosis, which implies that tumors diagnosed at different times arise from cells that carry the *SMARCB1-null or SMARCA4*-null genotype for very different periods of time (**Fig. 4e**). In 3 cases from SJD with available derived cell lines, we were able to approximate the calculation of the mutation rate to estimate the timing of the clonal expansion of each tumor. Although these results are an underestimation of the real timing –due to the unaccounted mutation rate resulting from cell culture passes–, these indicate that the MRCA clonal expansions occur at birth in one case, at 1 year old in the second case, and at age 7 in the third case. (**Extended Data Fig. 10**, see **Supplementary Note**–*Estimation of the chronological age of patients at the time of MRCA clonal expansions*). This opens the possibility that other non-genetic factors during the life of the child may be required to trigger the tumor expansion.

## Discussion

Here, we have exploited the genomic data of a large cohort of rhabdoid tumors to understand the timing of two key events in their evolution: the acquisition of their founder event –bi-allelic inactivation of *SMARCB1* or *SMARCA4*– and the most recent clonal expansion that gave rise to the malignancy. Our results indicate that the founder event of these tumors is very likely acquired during the children’s prenatal development. However, the most recent clonal expansion occurs at different moments along the life of the patients, relatively closer in time to the clinical diagnosis. Two recent studies on neuroblastoma, using the same rationale, also timed the main driver event (*MYCN* amplification) as very early, probably during the first trimester of pregnancy^20^ and the MRCA clonal expansion at different times (pre or postnatal) across cases^21^. Our study, thus, supports the widespread notion that many pediatric tumors, such as neuroblastoma and rhabdoid malignancies, have an embryonal origin^22^.

The results that place the acquisition of the driver bi-allelic inactivation early within prenatal development are robust, given the number of cases analyzed. Nevertheless, several limitations of the data and the approach prevent us from pinpointing a more accurate time. The limit of detection of somatic mutations imposed by the depth of sequencing and the purity, which determines that we focus on tumor clonal mutations, causes an overestimation of the time of acquisition of the founder event. Moreover, the two-speed model of the yearly age-related mutation rate, which accounts for differences during development and extra-uterine life, although useful for this purpose, is very simple and does not include potential differences between the embryonic (first trimester) and fetal (second and third trimester) development^19^. Finally, the mutation density data is noisy (as evidenced by the residuals of the regression), and that accounts for part of the variation in the estimated age of acquisition of the CN-LOH.

Our results match previous studies that suggested an embryonic origin of rhabdoid tumors. For example, work using mouse models showed that the time for *Smarcb1* inactivation dramatically affects the phenotype of the tumor, where early embryonic inactivation (early neural crest) of *Smarcb1* generates tumors that faithfully resemble human rhabdoid tumors, and later Smarcb1 inactivation leads to the development of schwannomas, a type of Schwann cells benign tumors^8,19^. These two types of tumors were also shown to be related in a separate study by Custers and colleagues, thus placing rhabdoid tumors on an ectodermal, neural crest-derived lineage, together with Schwann cells^10^. Another study on mouse models identified the potential cells of origin of MYC rhabdoid tumors, rooted in early stages of embryogenesis, as fetal primordial germ cells (PGCs)^9^. Moreover, our results –and those of other authors cited above^10^–indicate that cells that are *SMARCB1*-null or *SMARCA4*-null persist for some period of time –in some instances more than one decade– in the body without developing into a tumor. Phenotypically normal *SMARCB1*-null cells have indeed been shown to exist within healthy nerve sheath tissue in at least two cases^10^. Rhabdoid tumors, therefore, do not follow the *two-hit hypothesis* for tumor suppressor genes proposed by Alfred G. Knudson^23^, where the loss of the second allele immediately triggers tumorigenesis. Instead, a “third hit” –apparently non-genetic–is necessary for their development, similarly to other pediatric tumors^24^. For instance, in the case of Ewing sarcoma, some studies indicate that the first hit is an oncogenic fusion occurring during embryonic development and a second hit occurs at puberty with the increasing hormonal signals of IGF in the body, which trigger the development and growth of the cell harboring the oncogenic fusion^25^. In the case of rhabdoid tumors, although many are diagnosed at a very young age, it seems that another hit is required that promotes their development from a *SMARCB1*-null cell. Here, we borrow the concept of tumor promotion from adult cancer research^26^. An alternative explanation is the presence of anti-tumor mechanisms that keep in check the initiation of tumors in most cases, with the few that achieve immune evasion –taken from research in adult cancers^27^– resulting in tumorigenesis.

In the same line of the results obtained by Mel Greaves and colleagues in pediatric leukemia^24^, it is possible that *SMARCB1*-null or *SMARCA4*-null cells can be more widespread across children than the incidence of rhabdoid and other tumors. In agreement with our results, these cells would possibly emerge during embryonal development, a process that involves massive proliferation, apoptosis and rapid cell-fate shifting. The true incidence of *SMARCB1*-null or *SMARCA4*-null cells remains to be elucidated. The cause of the malignant transformation of these cells must also be an object of further research.

In conclusion, our work provides evidence that in human rhabdoid tumors the bi-allelic loss of *SMARCB1* or *SMARCA4* virtually always occurs very early during prenatal development, while the most recent clonal expansion of the tumor can occur many years later, always close to the age of diagnosis of the disease, probably promoted by other non-genetic factors.

## Methods

### Ethics approval

Patient material was included in the Sant Joan de Déu Barcelona Children’s Hospital Biobank for Research after informed consent from the patient’s legal guardians. Access to samples granted by the Hospital Sant Joan de Déu Drug Research Ethics Committee (CEIm) under approval number PIC-30-21. Solid biological material from the patient safeguarded in the Biobank was governed according to the Biomedical Research Law (Law 14/2007, of July 3, 2007), and by the provisions of Royal Decree 1716/2011, of November 18, 2011. The approval for the use of the Princess Maxima Cancer Center case was obtained from the Medical ethical committee of the Erasmus Medical Centre Rotterdam (the Netherlands). Informed written consent was provided by all patients and/or guardians.

### Sample collection, processing and biobanking

Tumor samples from SJD and a matched normal blood sample were collected at diagnosis unless specified. Fresh tumor samples containing at least 70% tumor cell content were immediately snap-frozen in liquid nitrogen or minced into small fragments (0.5 – 2 mm3) using sterile scalpels. The resulting tumor chunks were cryopreserved in fetal bovine serum (FBS) supplemented with 10% DMSO, or Synth-a-Freeze Cryopreservation medium (Gibco), and stored for long-term preservation in liquid nitrogen tanks for future use. MRT diagnosis was confirmed by immunohistochemical negative INI1 staining conducted by pathologists from the Anatomical Pathology Unit at the Sant Joan de Déu Children’s Hospital.

### DNA extraction

Samples from SJD were selected based on a pathologist-confirmed tumor purity of ≥ 70%, as assessed by H&E and INI1 negative stainings. Tumor genomic DNA was isolated using the QIAamp DNA Mini Kit (51304, QIAgen) according to the manufacturer’s instructions. DNA purity was assessed by measuring the A260/A280 and A260/A230 absorbance ratios using a NanoDrop™ 2000 Spectrophotometer (Thermo Fisher Scientific). Precise DNA concentration was further determined by fluorometric quantification using the Qubit™ dsDNA HS (High Sensitivity) assay kit on a Qubit™ X Fluorometer (Thermo Fisher Scientific). Library preparation was performed using KAPA HyperPrep Kits.

### *SMARCB1* mutation verification

Homozygous *SMARCB1* deletions from SJD samples were confirmed by multiplex ligation-dependent probe amplification (MLPA) using the SALSA MLPA Probemix P258 *SMARCB1* (P258-025R, MRC Holland); data analysis to confirm copy number variations were performed using Coffalyser.Net software (MRC Holland). For mutation verification, targeted PCR was performed to amplify the affected regions, followed by bidirectional Sanger sequencing (Macrogen, South Korea). Electropherograms were manually inspected and analyzed using Benchling (Benchling Inc., USA). Variants were identified by aligning the sequencing data against the *SMARCB1* reference sequence (NM_003073.5).

### Whole-genome DNA bulk sequencing and genomic analysis

Whole genome sequencing of SJD samples was performed from the tumor samples at a depth of 120X and the matched normal blood at a depth of 30X. Mutations were called using the matched normal samples as reference in each case.

### Collection of genomic data from repositories

Whole genomes of malignant rhabdoid tumor samples from the TARGET project were downloaded using the *Genomics Data Commons* (GDC) portal under an approved data transfer agreement through dbGaP (dbGaP Study Accession phs000218.v22.p8). Whole genomes of rhabdoid tumor samples from StJude were downloaded using the *St. Jude Cloud* (https://www.stjude.cloud) and DNANexus portals with the corresponding approved data transfer agreement^12^.

### Genomic analysis

The identification of somatic and germline variants was performed as previously described^28^. Briefly, after alignment of the tumor and normal genome reads, somatic single nucleotide variants (SNVs) and indels were identified using three calling tools (Mutect2^29^, Strelka2^30^, SAGE^31^ (https://github.com/hartwigmedical/hmftools/tree/master/sage)); Somatic Copy Number Variants and Structural Variants were identified using ASCAT^32^ and purple^33^. We further selected only variants identified within high quality mappable regions. To do this, a mappable genome was constructed as follows: 1) GEM Mapper tool^34^ was run to obtain the regions of hg38 genome with high degree of mappability; 2) blacklisted or problematic regions from ENCODE project were filtered out, using a file obtained from https://genome.ucsc.edu/cgi-bin/hgTables, on 14th May, 2025 ^35^. We finally filtered out common polymorphisms with allelic frequency above 0.001 across gnomAD 3.1 exomes and genomes^36^. The separation of clonal and subclonal mutations was carried out as described in^28^.

### Analysis of driver mutations

To identify driver alterations we focused on protein-affecting short variants (SNVs and indels) in each sample within genes included in the Compendium of Mutational Cancer Drivers Genes (at intOGen, release v2024; intogen.org). Our results only showed protein-affecting mutations in *SMARCB1* or *SMARCA4* genes. Therefore, we continued to further identify these mutations in the IGV bam file visualizer. Deletions involving the *SMARCB1* or *SMARCA4* genes identified by the mutation calling pipeline algorithm (purple^33^ and ASCAT^32^) were further inspected and visualized in the IGV bam file visualizer. On those samples where no driver event could be identified, *SMARCB1* and *SMARCA4* were further inspected in IGV bam file visualizer to identify any possible alteration not detected by the pipeline. In this detailed inspection, we were able to identify specific intra-genic deletions involving some exons.

### Mutational signature analysis

Mutational processes signatures were extracted using SigProfiler^37^. The tri-nucleotide change frequencies of all samples were generated using SigProfilerMatrixGenerator^38^ (version=1.3.4) and signatures were extracted, using all set of mutations (clonal and subclonal), for all the collected cohort using SigProfilerExtractor^39^ (version=1.2.1). Later, mutations were split into clonal and subclonal for each sample, and signatures were assigned using SigProfilerAssignment^40^ (version=0.2.3) with the set of mutational signatures identified for each sample in the extraction step.

### CN-LOH detection and mutation density calculation

CN-LOH events were detected using the B-allele frequency taken from the ASCAT Copy Number tool^32^. All genomic regions with two copies of the major allele and no copy of the minor allele, with a minimum length of 10Mb, were classified as CN-LOH events. We excluded samples harboring platinum treatment mutations (as per the signatures analysis). We filtered out SNVs that appear in the gnomADg database^36^ (version=3.1) with allele frequency > 0.001 (which are potentially polymorphisms) and mutations outside the boundaries of the mappable genome (see above). Samples with no mutations left in the CN-LOH region or with very aberrant copy number configurations in the chromosome with the CN-LOH were excluded. Only clonal mutations were selected for the timing analysis. The variant allele frequency (VAF) of each mutation was corrected by the tumor purity (cVAF = VAF / Purity). Mutations overlapping the CN-LOH region were then classified using this corrected VAF into homozygous (cVAF > 0.75) or heterozygous (cVAF < 0.75). Finally, the number of homozygous and heterozygous mutations were divided by the length of the CN-LOH region and multiplied by 10^6^ bases (N_mut_/length_CN-LOH_*10^6^ bases) to represent the density of homozygous (pre-CN-LOH) and heterozygous (post-CN-LOH) mutations per megabase in the CN-LOH. The mutation density in the entire genome of each tumor was computed using this same simple formula.

### Mutation rate calculation

We first computed the density of age-related mutations (selected using SigProfiler Assignment) along the entire tumor genome for each sample using the formula outlined in the previous section. An approximated yearly rate of the accumulation of age-related mutations in the genome of rhabdoid cells was calculated using the density of clonal age-related mutations per megabase of the treatment naive samples of the cohort, divided by the embryonic age of each sample (age at biopsy + 9 months of pregnancy). Then, a two-line regression model was defined between the mutation density and embryonic age at diagnosis in years. The first line in this regression follows the rate of accumulation of age-related mutations during prenatal development (0-9 months of embryonic age), while the second tracks the age-related mutation rate from birth to diagnosis. This two-line regression model is represented as follows:

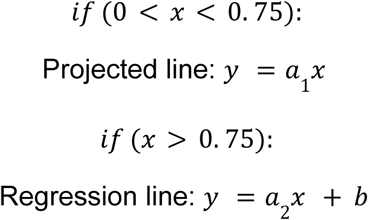

Where *x* is the embryonic age (in years), *y* is the number of mutations per megabase, *a*_1_ is the slope of the projected line from the zygote (0 years) to birth 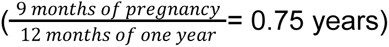, *a*_2_ is the slope of the regression line after birth, and *b* is the intercept (*y* value crossing at 0).

### Timing calculation

In order to calculate the chronological timing of the CN-LOH and the MRCA expansion of each sample, the two-line regression model was used to obtain the embryonic age (*x*) of each event by interpolating the CN-LOH homozygous (to time the CN-LOH) mutation density (*y*). Therefore, we apply the following formula:

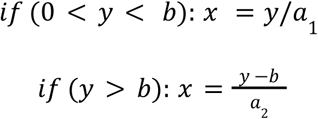

Further details on methodology can be found in **Supplementary Methods**.

## Data and code availability

Raw sequencing data (cram files) generated in this analysis for the samples of the 16 cases from SJD will be deposited in the European Genome-Phenome Archive (EGA) repository at the time of publication. All the code necessary to reproduce the figures of the paper will be made publicly available upon publication.

## Acknowledgements

N.L.-B. acknowledges funding from the European Research Council (consolidator grant 682398). M.S-G was supported by a postdoctoral contract with the Centro de Investigación Biomédica en Red de Cáncer (CIBERONC) and by the CRIS Emerging Leader grant (Emergingleader2025_04) from the CRIS Contra el Cáncer foundation. This project was supported by the CHEMOHEALTH project, funded by the Spanish Ministry of Science (MCIN), AEI /10.13039/501100011033/. It has also been supported by the project “Discovering the molecular signatures of cancer PROMotion to INform prevENTion” (PROMINENT) funded by Cancer Research UK (CGCATF-2021/100008), National Cancer Institute (1OT2CA278668-01) and the Spanish Cancer Association, AECC. IRB Barcelona is a recipient of a Severo Ochoa Centre of Excellence Award from the Spanish Ministry of Economy and Competitiveness (MINECO; Government of Spain) and an Excellence Institutional grant by the Asociacion Española contra el Cancer, and is supported by CERCA (Generalitat de Catalunya). We thank Ferran Muiños and Olivia Dove for their contribution in the discussion about the mutation rate calculation and the aging mutations regression. We are indebted to the “Biobanc de l’Hospital Infantil Sant Joan de Déu per a la Investigació” integrated in the Spanish Biobank Network of ISCIII for the sample and data procurement. We are grateful to the Band of Parents at Hospital Sant Joan de Déu Barcelona for supporting the overall research activities of the developmental tumor laboratory, PCCB, and for having generously contributed with their samples. This study would have been impossible to reach without this contribution.

## Notes

### Competing Interest Statement

The authors have declared no competing interest.

