## Supplementary information for "Variable latency between the founder genetic event and rhabdoid tumor expansion"

**Affiliations**

**Supplementary Methods** **3**
Mislabeled neuroblastoma sample 3
Clonal expansions of single cells from cell lines derived from a tumor sample 3
**Supplementary Note** **4**
A case with two germline *SMARCB1* variants 4
Evolution of multiple rhabdoid tumors in an individual 4
Estimation of the chronological age of patients at the time of MRCA clonal expansions 4
**Extended Data Figures** **6**
**References** **19**

### Supplementary Methods

#### Mislabeled neuroblastoma sample

All samples harbored loss-of-function mutations that targeted both alleles of either *SMARCB1* or *SMARCA4* genes, except for the PT51 sample of the TARGET cohort. In this case, a *MYCN* amplification was found to be the main driver event. We further explored this sample to understand if this is a true rhabdoid tumor by the analysis of the methylation profile of this sample[1] and applied the pediatric tumor classification based on methylation to it. Briefly, the methylation data for the tumor of this patient, obtained via the Infinium Methylation EPIC v2.0 BeadChip (Illumina) was run through the Heidelberg CNS Tumor Methylation Classifier v12.8 available at https://app.epignostix.com for methylation-based classification and chromosomal copy number (CNV) analysis based on the raw intensities of the methylation array probes. The results indicated that this tumor is a neuroblastoma with a 97% accuracy (**Extended Data Fig. 3a)**. In agreement with this, the mutational profile of this sample showed higher frequency of C>A substitutions, which are characteristic of the activity of SBS18 signature, very active in neuroblastomas[2] (**Extended Data Fig. 3b**). We therefore decided to exclude this sample from the analyses of this cohort.

#### Clonal expansions of single cells from cell lines derived from a tumor sample

Patient-derived rhabdoid tumor cell lines were obtained from a small fraction of fresh or cryopreserved tumor fragments. For derivation of the PT84 cell line, an *in vivo* amplification step in athymic nude mice was required prior to in vitro culture. PT84 xenograft tumors were subsequently harvested and processed for *in vitro* culture. Tumor fragments from PT53, PT81 and PT84 were enzymatically digested in RPMI 1640 containing 200 U/mL collagenase (Sigma, C5138), 160 U/mL DNase I (Sigma, D4263) for 30 min at 37ºC under gentle agitation. Following a wash step, the digested samples were further dissociated with 0.25% Trypsin-EDTA (Gibco, 25200056) at 37 ºC for 5 min. After trypsin inactivation, the cell suspension was filtered through a 100-µm cell strainer and equal aliquots were cultured in parallel in three different media to determine the most suitable conditions. Based on these results, PT53 cell line was maintained in DMEM/F12 medium (Gibco, 21331020) supplemented with 1% B27 supplement (Gibco, 17504044), 20 ng/mL Human EGF Recombinant protein (Gibco, PHG0311), 20 ng/mL Human FGF-basic Recombinant Protein (Gibco, PHG0026) and 100 U/mL penicillin-streptomycin (Gibco, 15140122). PT81 and PT84 cell lines were cultured in TSMserum medium consisting of 50/50 Neurobasal A medium (Gibco, 10888022) and DMEM/F12 (Gibco, 21331020), supplemented with 10 mM HEPES buffer (Gibco, 15630080), 1 mM Sodium-pyruvate (Gibco, 11360039), 0,1 mM MEM non-essential amino acid solution (Gibco, 11140035), 1% Glutamax-1 supplement (Gibco, 35050061), 100 U/mL penicillin-streptomycin (Gibco, 15140122) and 10% FBS (Gibco, A5256701). Fresh growth media was added every 48 to 72 hours, and cultures were maintained at 37ºC in a humidified incubator at 5% $CO_2$. Absence of mycoplasma contamination was routinely tested using the EZ-PCR Mycoplasma Detection kit (Biological Industries, 20-700-20) according to the manufacturer's instructions.

All cell lines were sorted and expanded to obtain single cell clonal expansion. PT53 cell line was sorted at passage 12 (P12), PT81 cell line was sorted at passage 10 (P10) and PT84 cell line was sorted at passage 5 (P5). For each cell line, two clones were selected and expanded to collect pellets for gDNA isolation (as previously described).

### Supplementary Note

#### A case with two germline *SMARCB1* variants

We explored the intriguing PT76 case, where we identified two germline *SMARCB1* alterations. We annotated the location and the protein impact of both mutations: one is a frameshift indel affecting the first exon (F10X) and the other is a nonsense SNV located in the fifth exon (W206*) (**Extended Data Fig. 3c**). The two variants are present in both the tumor and the blood samples, at an allele frequency of ~0.5, pointing to a likely germline (or very early somatic mosaic) origin of the two mutations (**Extended Data Fig. 3d**). We are not aware of any similar case described in the scientific literature. We speculate that some *SMARCB1* protein activity may be retained by one of the two alleles; for example, there could be an alternative transcript skipping the 1st or the 5th exon resulting in a partially functional *SMARCB1* protein.

Furthermore, the mutational profile of this sample harbored an unusual number of T>G mutations (**Extended Data Fig. 3e**), which was already described in the previous publication from the TARGET cohort[3]. This could be explained by a high relative activity of SBS17a and SBS17b mutational signatures across the cells of the sample (**Extended Data Fig. 4a**). SBS17b has been found almost exclusively in samples of the digestive tract, mostly esophagus and stomach[4,5], or from other tissues in individuals exposed to 5-fluorouracil/capecitabine treatment[6]. The activity of these mutational signatures could be providing a clue as to the cell of origin of this tumor. However, according to the clinical data from the TARGET project, it is located in the kidney. Of note, another case showing SBS17a and SBS17b (PT86) is located in the paraspinal area. Whether any of these tumors indeed originated from a cell developmentally related with the gastrointestinal tract remains to be investigated.

#### Evolution of multiple rhabdoid tumors in an individual

The evolution analysis of cases with two or more samples show that rhabdoid tumors are of rapid expansion and dissemination, especially in younger children. In older children (PT83 and PT86, with eight and thirteen years respectively), a higher number of unique clonal mutations (i.e., accumulated after the separation of the rhabdoid lineages giving rise to the tumors) in each tumor was observed. This likely points to an earlier time of divergence of these rhabdoid lineages. The spread of the tumors in these cases occurred prior to the diagnosis of the first malignancy.

An interesting case is PT81 from SJD. This case represents a patient diagnosed with a MRT of the kidney at 4 years and 9 months old, who 2 years and 4 months later relapsed with a metastatic lesion in the lung. Here, almost a third of the clonal mutations unique to the metastatic tumor were contributed by carboplatin. Interestingly, this case showed a much higher number of aging mutations in the relapse tumor, possibly due to an accelerated aging due to the exposure to chemotherapy. In fact, some recent studies point to the possibility that the activity of SBS5 could be increased by exogenous DNA damage agents[7] (**Extended Data Fig. 5b**).

We further analysed the case of PT70, which presents an interesting example of tumor evolution. Four samples of four different malignant lesions were obtained from PMC at autopsy. The evolutionary analysis shows that likely 3 separate tumor expansions took place in this patient. The first clonal expansion gave rise to T1 (affecting a lymph node); the second gave rise to T2 (affecting a separate lymph node) and T3 (lodged in the omentum); the third clonal expansion originated T4 (in the diaphragm) (**Extended Data Fig. 5c**).

#### Estimation of the chronological age of patients at the time of MRCA clonal expansions

To provide a separate estimation of the chronological age of individuals at which the MRCA of the tumor experienced the latest clonal expansion, we used cell lines derived from tumor samples from the tumors of cases PT53, PT81 and PT84 (all from the SJD cohort). We reasoned that the total number of mutations (clonal, subclonal and private SNVs) in the tumor cell expanded to form the cell line will be expanded, and could therefore be detected via whole-genome sequencing of the cell culture (**Extended Data Fig. 10a**). All clonal mutations detected in this culture were, thus, present in a single cell of the rhabdoid tumor and had accumulated between the first cell division of the zygote and the moment of the biopsy. These could then directly be used to determine the rate of accumulation of age-related mutations in cells giving rise to the rhabdoid tumors in these donors. For robustness, we whole-genome sequenced two separated cell expansions obtained from each tumor (**Extended Data Fig. 10b**).

However, one important limitation of this assay is that the cell lines available have been passed a number of times before the establishment of the definitive cell line, which we could sort into single cells. In these passages, a number of age-related mutations have accumulated which will result in an overestimation of their rate of accumulation (**Extended Data Fig. 10c**). Indeed, these mutation rate values are higher than those estimated using only age-related clonal tumor mutations (explained in the main manuscript) (**Extended Data Fig. 10d**). Therefore, the age of the most recent rhabdoid clonal expansion is underestimated using the mutation rate values calculated from these single cell expansions. As a result, the true most recent rhabdoid clonal expansion will have taken place later than estimated.

Having only three cases with available cell lines, we decided to perform a very simplistic analysis per case. We directly calculated the yearly rate of accumulation of age-related mutations in the rhabdoid lineage as the number of clonal age-related mutations identified in the whole-genome of a single cell expansion divided by the age of the donor at the time of biopsy. Then, we divided the number of clonal mutations detected in the tumor (from which the cell line was derived) at the time of biopsy by this rate, thus obtaining an estimate of the time that explains that accumulation. As these ages are underestimated (see above), they provide a lower limit for the true age of the last clonal expansion. In case PT53 we can estimate that  the expansion that gave rise to the MRCA occurred, at the earliest, right after birth. In case PT81, the MRCA of T1 expanded not earlier than 1 year of age, and the MRCA of T2 expanded, at the earliest, by 3 years and 9 months of age. Finally, for PT84, the MRCA of T1 expanded, at the earliest, by 7 years of age. Interestingly, this case (PT84) was diagnosed with a neuroblastoma at 1 year and 5 months old and 8 years later with a rhabdoid tumor (at 9 years and 11 months old)[8]. While in a previous article describing this case we were unable to time the expansion of the MRCA of the rhabdoid, here we are able to do it through this methodology. As we estimate that the MRCA of this tumor expanded not earlier than 7 years of age, this expansion occurred 6 years or more after the diagnosis of the neuroblastoma.

We are unable, in these cases, to estimate the age of emergence of the *SMARCB1* null cell, since only the case PT53 harbors a CN-LOH event; unfortunately, no homozygous mutation overlaps the genomic region of the event. This makes it impossible to estimate this age.

This analysis is a first attempt to estimate the age of occurrence of the MRCA clonal expansion that ultimately gave rise to the rhabdoid tumors in these patients. Future analyses are needed to refine these calculations. However, these results already indicate that the MRCA clonal expansion is, in all likelihood, postnatal, while the acquisition of the *SMARCB1/SMARCA4* null cell is very likely prenatal.

### Extended Data Figures

#### Extended Data Figure 1

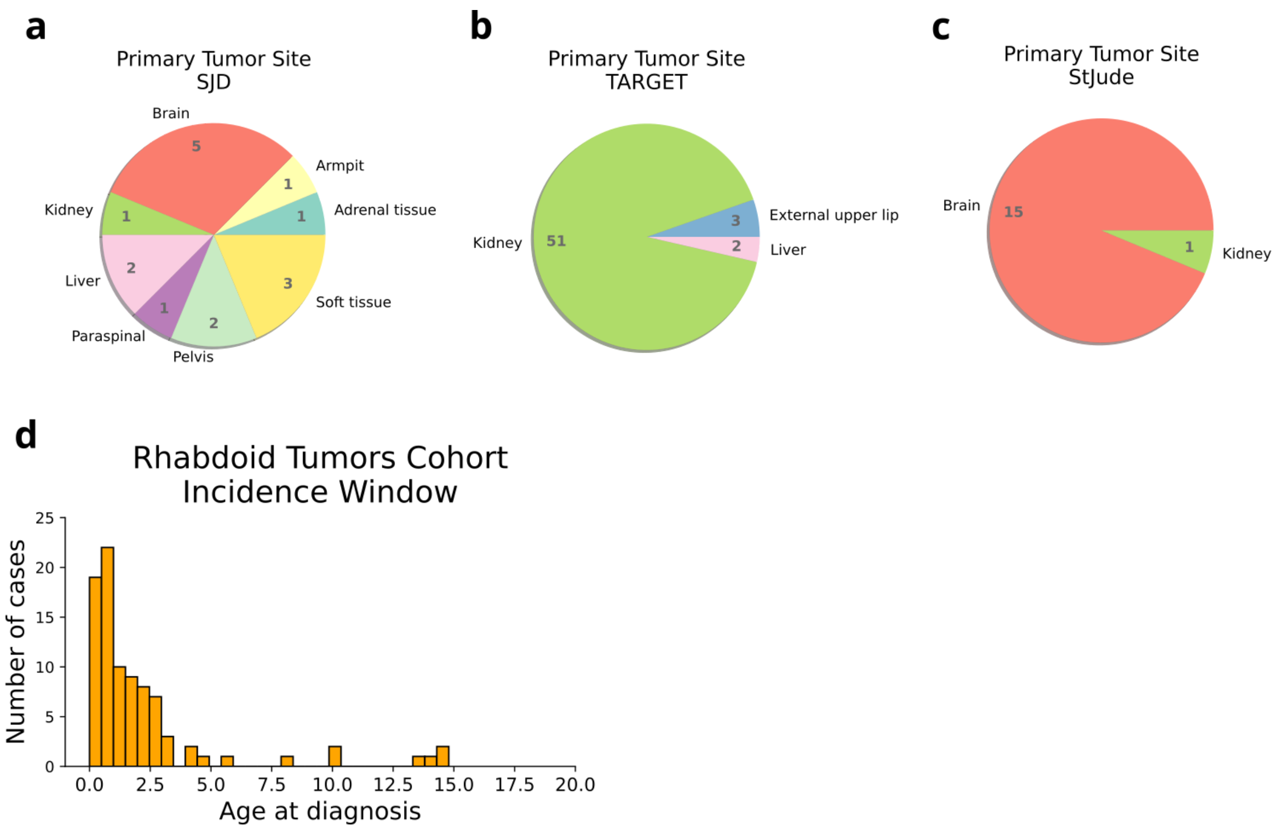


**Extended Data Figure 1. Extended clinical overview of the rhabdoid tumors cohort.**
**a**, Primary tumor site of SJD; **b**, Primary tumor site of TARGET; **c**, Primary tumor site of St Jude; **d**, Incidence age window for the whole rhabdoid tumor cohort.

**Extended Data Figure 2**

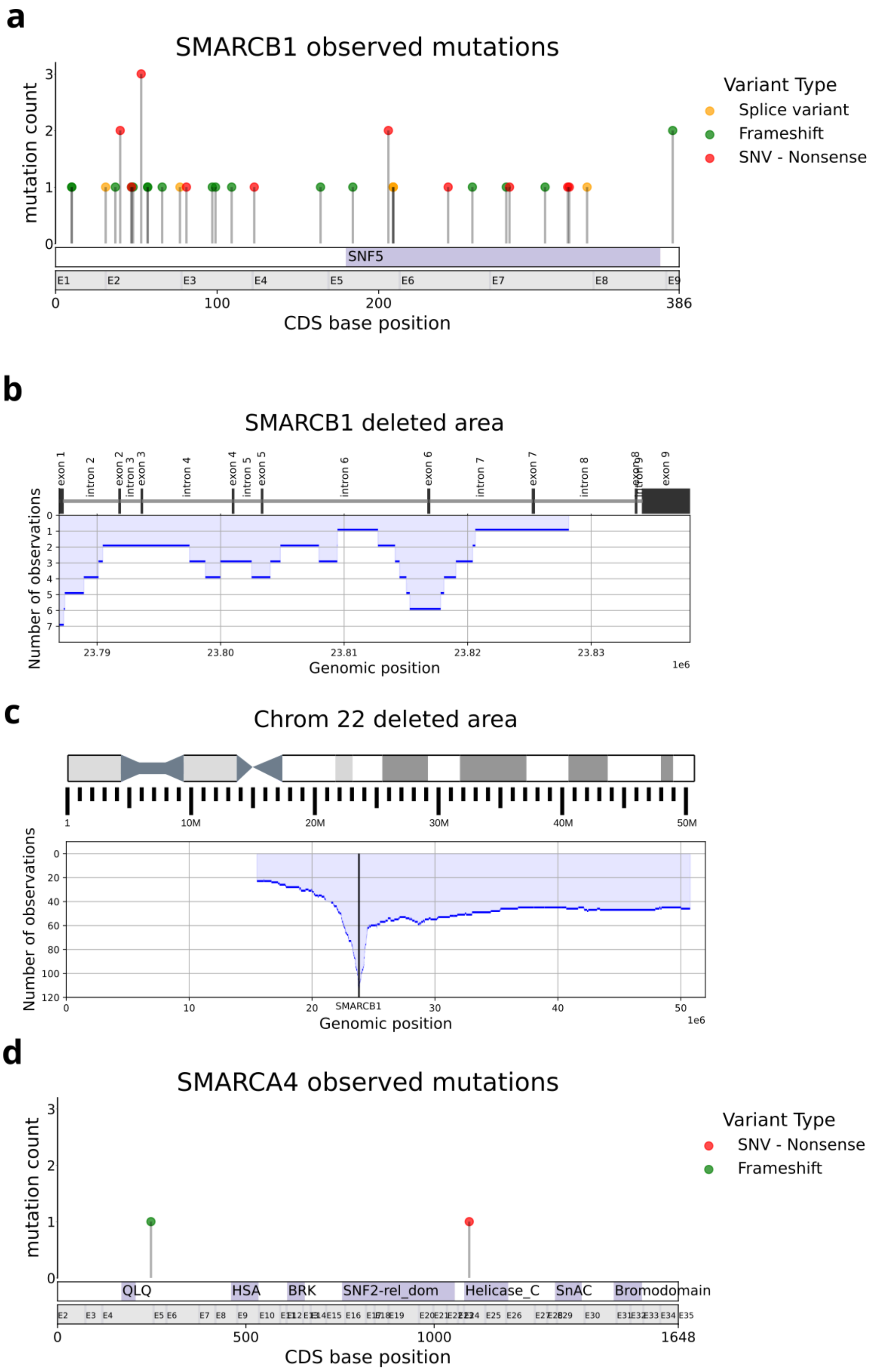


**Extended Data Fig. 2**: ***SMARCB1* and *SMARCA4* alterations overview.**

**a**, Distribution of SNV and INDEL mutations found in *SMARCB1* gene; **b**, Distribution of exonic deletions in *SMARCB1* gene; **c**, Distribution of the deleted area (deep deletions) of chromosome 22; **d**, Distribution of snv and indel mutations found in *SMARCA4* gene; **e**, Distribution of the deleted area (deep deletions) of chromosome 19.

### Extended Data Figure 3

**a**

**Methylation classification**

| Predictions | | | | Calibrated score | Interpretation | |
|---|---|---|---|---|---|---|
| neuroblastoma | | | | 0.97962 | match | ✔ |
| | neuroblastoma | | | 0.97962 | match | ✔ |
| | | neuroblastoma | | 0.97962 | match | ✔ |
| | | | MC Neuroblastoma, MYCN type | 0.97514 | match | ✔ |

Legend*: ✔ Match (score >= 0.9) ✘ No match (score < 0.9): possibly still relevant for cases with low tumor content and poor DNA quality.

Hierarchy levels:

| | | | |
|---|---|---|---|
| Superfamily | | | |
| | Family | | |
| | | Class | |
| | | | Subclass |

*Only predictions with scores >= 0.9 are considered "classified". Scores < 0.9 could still be relevant for cases with low tumor content or poor DNA quality. Please exercise caution in interpreting such predictions.
Only superfamily predictions with scores >=0.3 are shown, lower hierarchical levels are displayed if they are predicted with scores >=0.1

***Description:***

MC Neuroblastoma, MYCN type: The "mc Neuroblastoma, MYCN subtype" represents a subgroup of neuroblastoma - a peripheral neuroblastic tumour with a neural crest origin arising in childhood. The tumours of this mc often show activation of MYCN, commonly through genetic amplification. This mc refers to 'typical' peripheral neuroblastoma, as distinct from "mc CNS neuroblastoma, FOXR2-activated".
Evidence Level: None

**b**

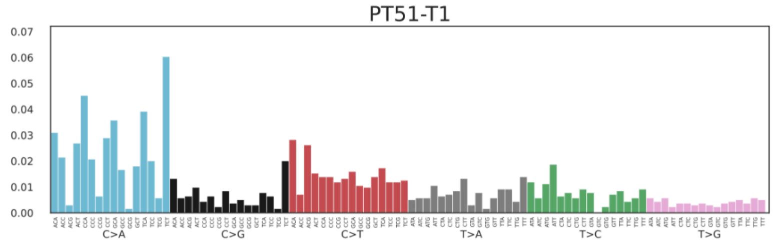


**c**

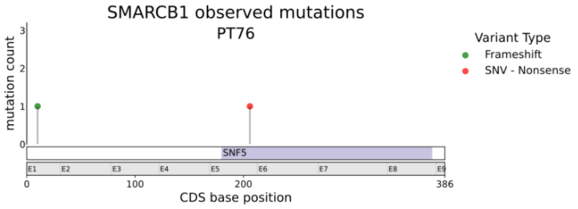


**d**

| SNVs/Indels | | | | Tumour sample | | | | Normal (Blood) sample | | | |
|---|---|---|---|---|---|---|---|---|---|---|---|
| variant_type | Consequence | aa_change | mut | AF | N alt reads | N ref reads | Depth | AF | N alt reads | N ref reads | Depth |
| truncating | stop_gained | W206* | chr22:23803411:G>A | 0.62 | 18 | 17 | 35 | 0.34 | 10 | 19 | 29 |
| truncating | frameshift_variant | F10X | chr22:23787197:TT>- | 0.24 | 7 | 15 | 22 | 0.55 | 16 | 11 | 27 |

**e**

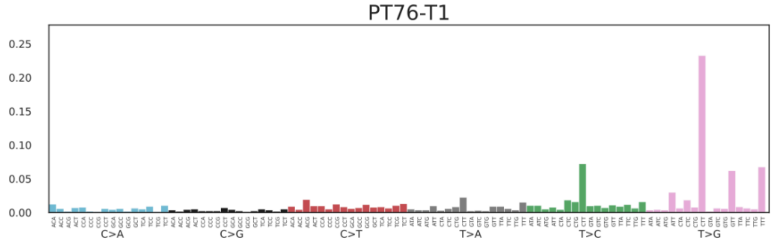


**Extended Data Figure 3: PT51 case classifies as a neuroblastoma and PT76 case with two germline mutations.**

**a**, Methylation profile analysis of PT51 showing the tumor classification as a neuroblastoma; **b**, Mutational profile of PT51 sample; **c**, Needle plot of PT76 sample showing the 2 germline mutations; **d**, Table with detailed information about the two detected germline mutations in PT76; **e**, Mutational profile of PT76 sample.

**Extended Data Figure 4**

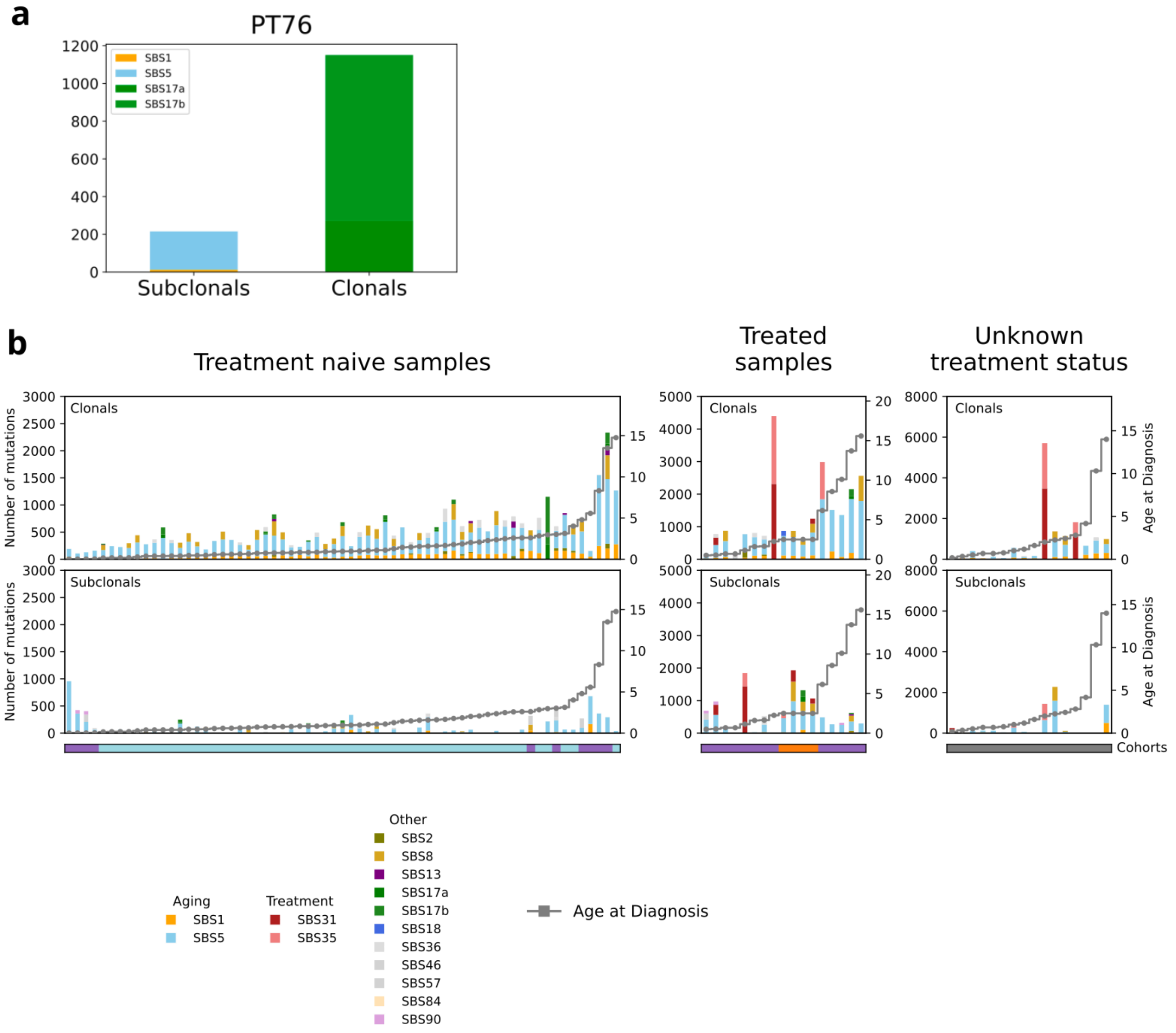


**Extended Data Figure 4: Mutational signatures assignment per sample for clonal and subclonal settings**

**a**, Clonal and subclonal mutational signatures assignment for PT76 case; **b**, Clonal and subclonal mutational signatures assignment for all the rhabdoid tumors cohort.

### Extended Data Figure 5

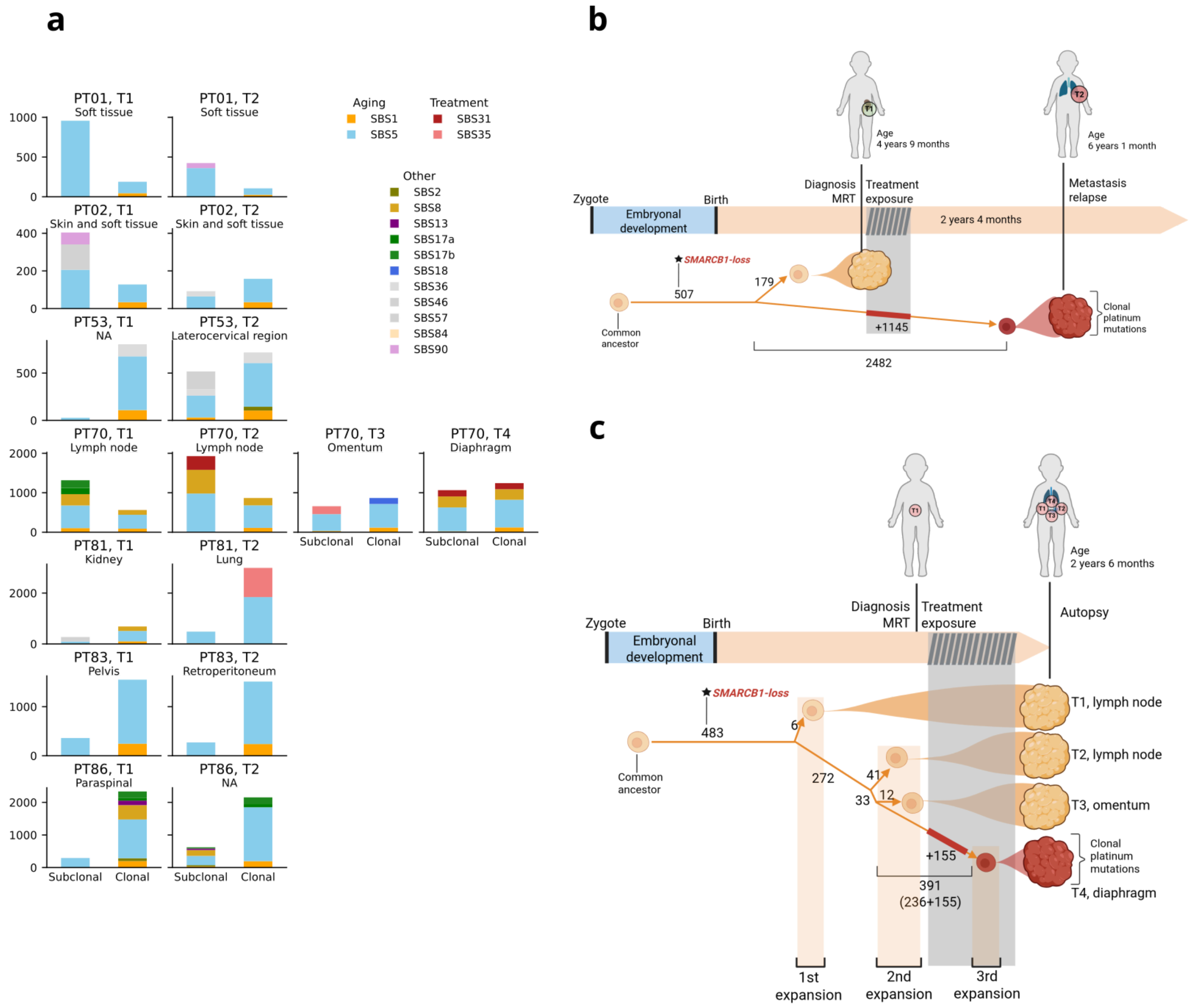


### Extended Data Figure 5: Extended analysis on cases with several tumor lesions.

**a**, Number of mutations of clonal and subclonal settings of each tumor sample of each case, indicating the fraction of mutations that correspond to each mutational signature; **b**, Tumor evolution of PT81; **c**, Tumor evolution of PT70.

**Extended Data Figure 6**

**a**

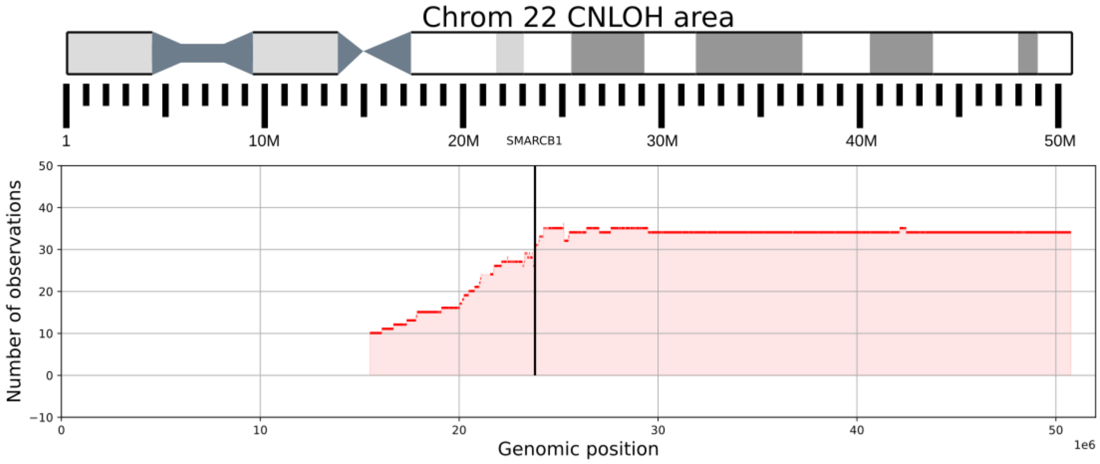


**b**

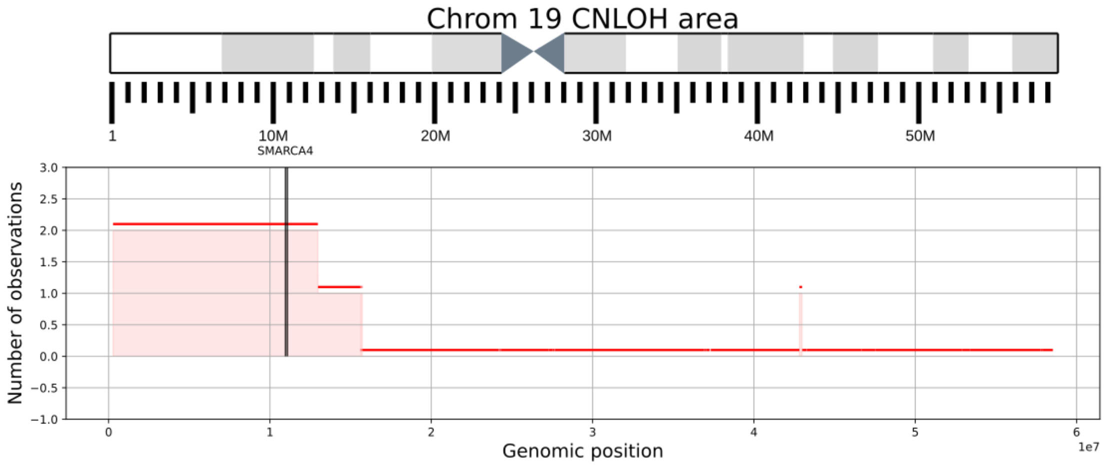


**Extended Data Figure 6. Chromosomal area covered by the CN-LOH.**

**a**, Distribution of the CN-LOH of chromosome 22; **b**, Distribution of the CN-LOH of chromosome 19.

### Extended Data Figure 7

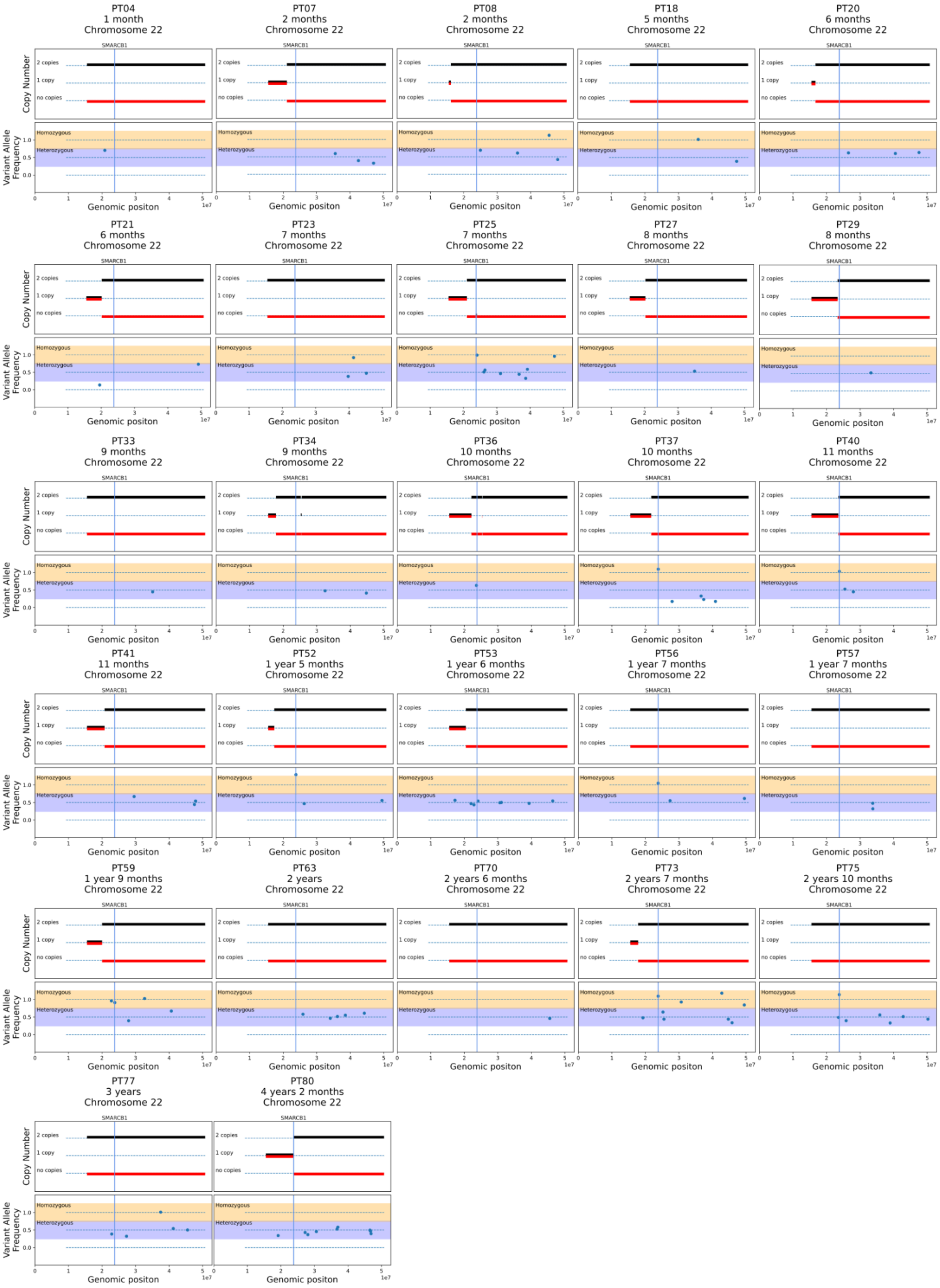


**Extended Data Figure 7**: **Distribution of mutations in the CN-LOH area for each case.**

Gained chromosome copy is represented in black and lost chromosome copy is represented in red. Homozygous mutations are located in the orange area, and heterozygous mutations are located in the blue area.

**Extended Data Figure 8**

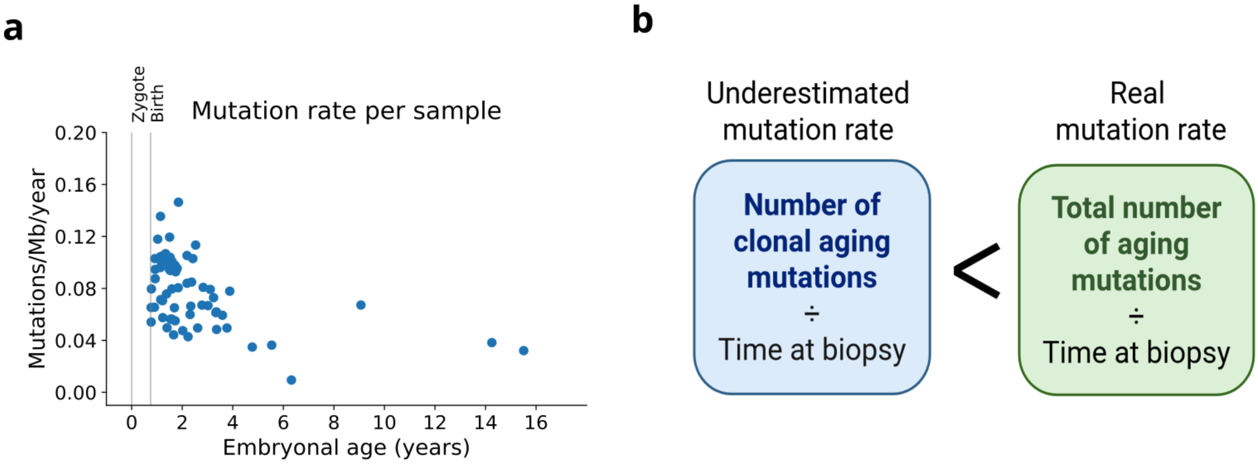


**Extended Data Figure 8. Approximation of mutation rate with the clonal aging mutations**.

**a**, Distribution of mutation rate (mutations per megabase per year) across the embryonic age (age at diagnosis + 9 months of pregnancy) for each treatment naive sample; calculated using the clonal aging mutations; **b**, The mutation rate calculated with the clonal aging mutations is an underestimated mutation rate in respect to the real mutation rate.

**Extended Data Figure 9**

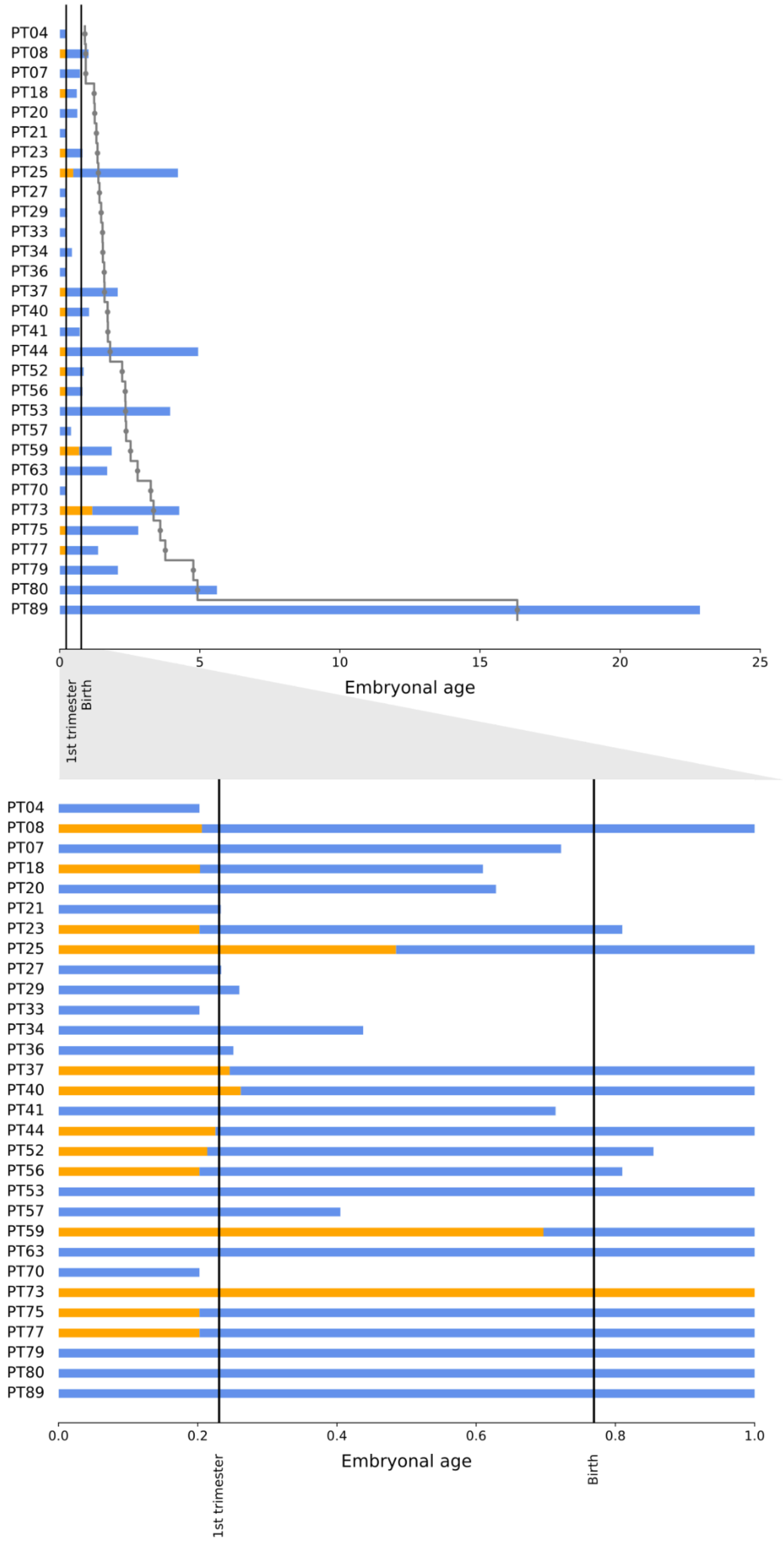


**Extended Data Figure 9**. **Timing the CN-LOH across cases.**

Orange bars represent time until CN-LOH and blue bars indicate time until MRCA. Top panel: Age range from 0 to 25 embryonal age represented. Age at diagnosis is indicated in grey; Bottom panel: Age range from 0 to 1 embryonal age represented.

### Extended Data Figure 10

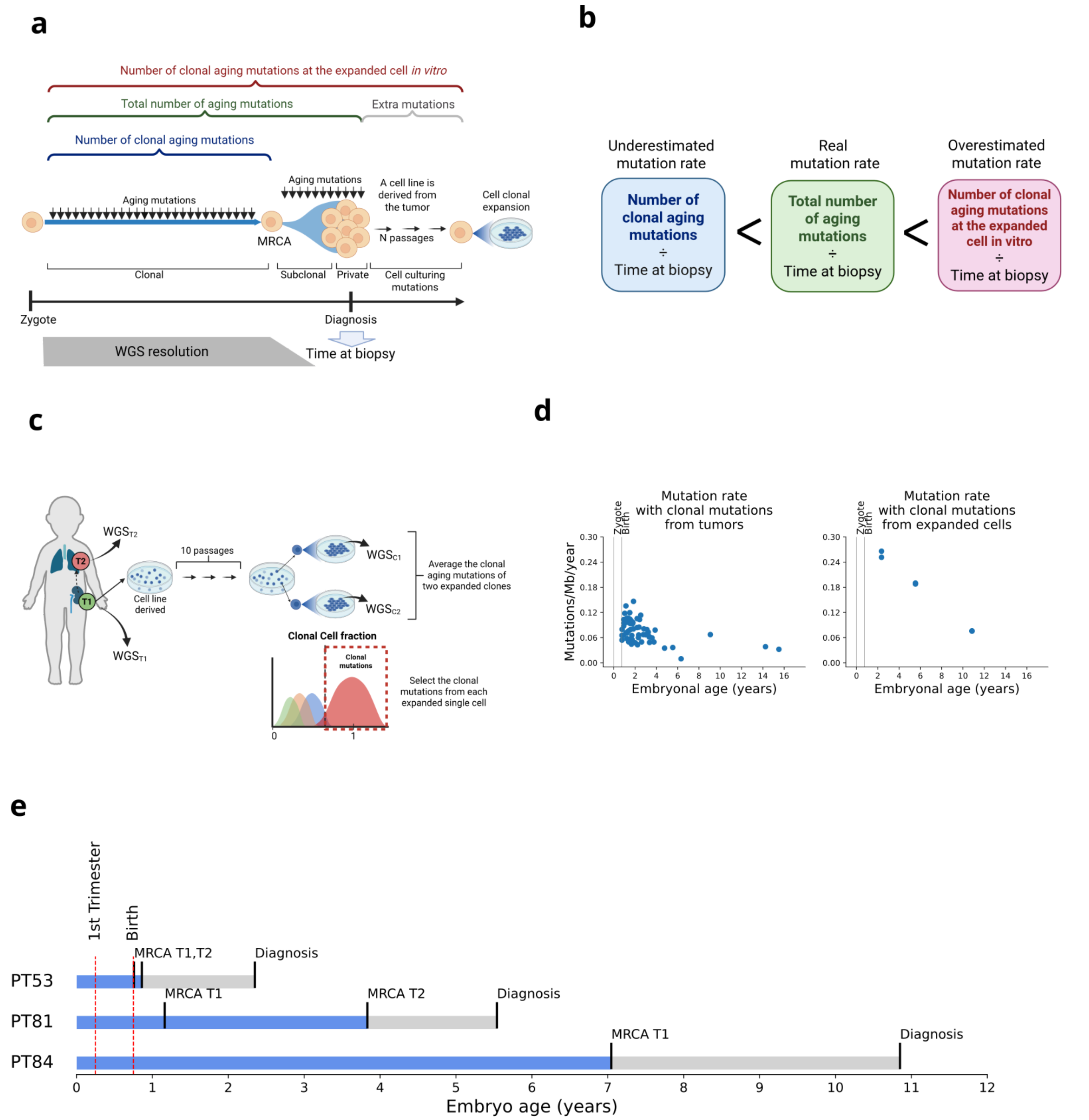


**Extended Data Figure 10**: **Timing of the MRCA clonal expansion in 3 cases with derived cell lines, using clonal aging mutations of expanded cell clones**.

**a**, Representation of the mutations detected in the in vitro expanded clones, indicating that extra mutations are accumulated in the passages between the tumor collection and the single cell clonal expansion; **b**, Representation of PT81 sample collection and cell line derivation from the kidney-located tumor sample. The expanded single cells come from an aliquot with 10 passages from the cell line derivation. The clonal mutations of the expanded single cells represent the total number of mutations accumulated until the time of the biopsy plus the extra mutations accumulated during cell culturing; **c**, The mutation rate calculated with the clonal aging mutations at the expanded cell *in vitro* is overestimated due to the extra mutations added during cell culture; **d**, Mutation rate (mutations per megabase per year) per case (left) and mutation rate (mutations per megabase per year) per expanded cell in vitro (right); **e**, Timing of MRCA per case. Blue bar indicates time from the zygote to the MRCA clonal expansion of the tumors (T1 and/or T2). Grey bars indicate the time from the MRCA clonal expansion of the latest tumor to the time of diagnosis. Indicated in red are the end of the 1st trimester and the time of birth.
